# ScriptManager: a platform for scalable and reproducible high-resolution analysis of genomics datasets

**DOI:** 10.64898/2026.06.14.732163

**Authors:** Olivia W Lang, Benjamin Beer, Dean Zhang, Courtney LeSon, Alima Deen, B. Franklin Pugh, William KM Lai

**Affiliations:** Cornell EpiGenomics Facility, Cornell Institute of Biotechnology, Cornell University, Ithaca, New York, United States of America; Department of Molecular Biology and Genetics, Cornell University, Ithaca, NY 14853, USA; Department of Computational Biology, Cornell University, Ithaca, NY 14853, USA

**Keywords:** Epigenomics, Genomics, FAIR, Chromatin, ChIP-exo, ATAC-seq, PRO-cap, High Throughput Computing

## Abstract

**Background:** The growing diversity of genomic and epigenomic assays has driven a parallel expansion in data formats, analysis workflows, and figure-generation tools. However, tools for analyzing data and assembling publication-quality figures are often specialized to a specific assay, dramatically limiting their interoperability and reproducibility.

**Results:** We present the v1.0 release of ScriptManager, a Java-based framework for modular and reproducible analysis and visualization workflows of genomics and epigenomics data. Unlike existing tools specialized for individual assay types, ScriptManager provides a unified and extensible framework for cross-assay visualization and workflow reproducibility. The v1.0 release adds novel analytical modules, GUI session logging, automated unit and integration testing, tutorials, and expanded documentation. It also integrates with the broader reproducibility ecosystem through Singularity containers, Anaconda packaging, and Galaxy XML wrappers. We demonstrate ScriptManager’s TagPileup scaling from local single-core execution to a 10,305-job analysis distributed across the Open Science Grid (OSG), with the full workload completing in <2 hours of wall-clock time.

**Conclusions:** ScriptManager v1.0 enhances workflow portability, transparency, and reproducibility across a diverse range of high-resolution genomic assays. By coupling a flexible module design with modern reproducibility standards, ScriptManager provides a bridge between exploratory data analysis and formal, publication-ready figure generation. These improvements enable researchers to build, share, and reproduce genomic analyses across diverse computational infrastructures with minimal configuration.

## Background

The last two decades of genomics technology development have seen an explosion of novel biochemical assays that leverage high-throughput sequencing. Foundational assays like RNA-seq have branched into sub-types that capture different subpopulations of transcripts (e.g., CAGE, PRO-seq) while others profile chromatin architecture and accessibility (e.g., ATAC-seq, CUT&Tag, ChIP-exo) [1, 2]. While these assays have enabled genome-wide characterization of transcriptional activity, chromatin accessibility, and protein-DNA interactions at unprecedented resolution, their corresponding analysis tools remain highly specialized. This has resulted in an increasingly fragmented software landscape where a given assay’s analysis and visualization toolkits are rarely portable across different assays or compute environments.

This fragmentation has amplified challenges in reproducibility and workflow maintenance. Bioinformatic tools range from general utilities (e.g., samtools, bedtools, FastQC) to structured end-to-end pipelines (e.g., SLAM-Dunk, GATK) [3–9]. Modular architectures offer flexibility but can be difficult to reproduce and document, while monolithic pipelines provide stability at the expense of user control [10]. Balancing these philosophies while meeting increasingly stringent Findable, Accessible, Interoperable, and Reproducible (FAIR) standards remains a central challenge [11]. Even when code is shared, incomplete documentation of parameters, software versions, or computational environments often prevents full replication, and differences in operating systems, dependency versions, or resource configurations compound these failures [12–14].

Reproducibility frameworks, including workflow managers (e.g., Galaxy, Nextflow, Pegasus), dependency isolation systems (e.g., Anaconda), and containerization technologies (e.g., Docker, Singularity) have emerged to address these barriers [15–20]. However, adoption of these frameworks remains difficult for small laboratories and interdisciplinary users who lack dedicated computational support. Concurrently, funding agencies and journals are now enforcing rigorous reproducibility standards [21, 22]. The challenge is especially acute during exploratory data analysis. Researchers typically iterate rapidly through parameter combinations and visualization strategies that often form the basis of publication figures. These parameters are rarely recorded with sufficient detail for others to reproduce [23, 24]. Critically, without automated mechanisms to capture parameters, dependencies, and data provenance, the transition from *ad hoc* exploratory scripts to formal, shareable workflows results in loss of critical information [10].

To help close the gap between exploratory flexibility and formal reproducibility, we developed ScriptManager, a modular Java-based framework for genomic data analysis and visualization. ScriptManager provides a structured yet extensible environment for executing analysis scripts across diverse assay types. We previously presented a proof-of-concept of ScriptManager as a platform for both GUI-based and command-line genomic analysis [25]. Here we present the v1.0 release: a fully featured and scalable platform that supports log-based reproducibility, Open Science Grid-powered scalability, comprehensive tutorials, and an expanded tool suite for a wide variety of genomic and transcriptomic assays. These updates collectively establish ScriptManager as a robust, FAIR-compliant platform for reproducible genomic workflow construction and visualization.

## Implementation

ScriptManager’s design and architecture, including its graphical and command line interfaces, were designed for ease-of-use and convenience for both the user and future development. Beyond its dual GUI/CLI design, ScriptManager v1.0 introduces several features specifically aimed at reproducibility, including session logging, stochastic seeding, containerization, and unit testing.

### Design and Architecture

ScriptManager is a suite of genomic analysis tools implemented in Java that can be executed through either a graphical user interface (GUI) or a command line interface (CLI). The suite includes novel analytical implementations, re-implementations of existing methodologies, and GUI wrappers for common Java-based command-line tools (i.e., Picard) (**Fig. 1A**, **Additional File 1**) [26]. ScriptManager v1.0 includes 47 total tools, representing a dramatic expansion in tool capabilities and platform features since the original proof-of-concept [25].

**Figure 1:**
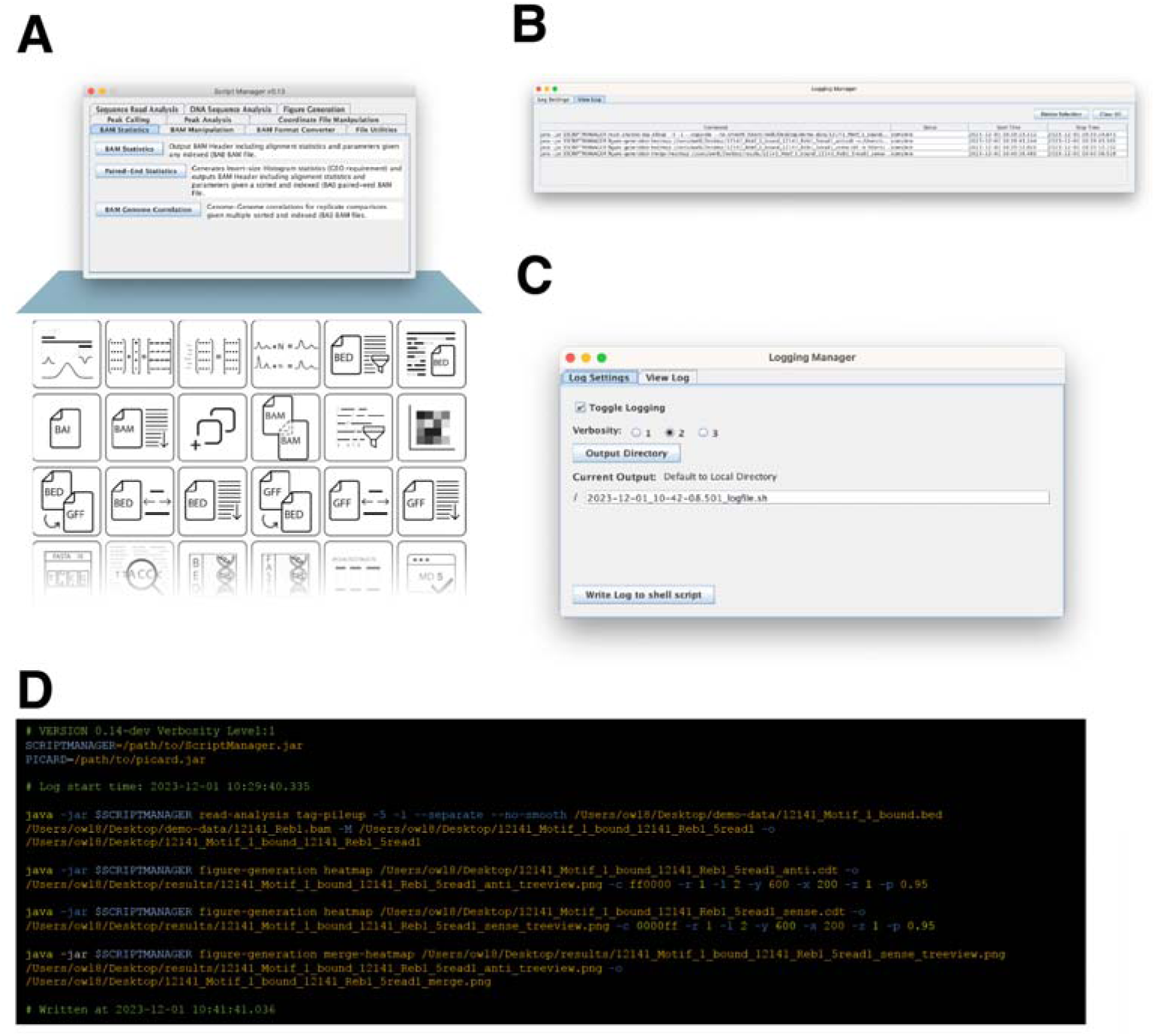
ScriptManager v1.0 includes logging support for the entire collection of tools. (A) Any script in the collection can be launched by selecting it from the main window. The Logging Manager window can also be accessed from the main window. (B) Opening the manager allows the user to view the entire log under the “View log” tab of the log manager where entries are updated with an execution status of “incomplete”, “error”, or “complete.” (C) Under the “Log Settings” tab the user can toggle logging on/off and set options for saving the log. (D) The saved log is a shell script of each script execution with all option information in the GUI mapped to the ScriptManager CLI execution statements.

A Java-based implementation was chosen to balance performance, portability, accessibility, and software longevity. ScriptManager leverages the HTSJDK library for efficient parsing of genomic file formats and supports multithreaded execution for computationally intensive operations, enabling it to handle the large datasets common in genomics [26]. The application is compiled as a single executable JAR that contains all dependencies needed for analysis and visualization, bypassing the dependency installation failures, version clashes, and cross-platform incompatibilities that frequently impede adoption of bioinformatics software [14].

To ensure interoperability with the broader genomic tool ecosystem, ScriptManager operates on widely adopted standard file formats including BAM, BED, GFF, and customizable tab-delimited matrix formats (.cdt, .tsv, .txt). This adherence to community standards means that users can freely combine ScriptManager tools with third-party software in a single workflow: when an analysis step falls outside ScriptManager’s scope, users can hand off to an external tool and return to ScriptManager without format conversion, provided the external tool also follows standard genomic file conventions. ScriptManager’s documentation includes lookup tables that index tools by their expected input and output formats to help users identify appropriate tools and plan interoperable workflows.

### User Interfaces – GUI & CLI

Researchers new to bioinformatics (or to any new software) typically face a learning curve that can delay their first analytical results as they learn to navigate new systems and commands. ScriptManager includes a Swing-based graphical user interface (GUI) designed to guide novice bioinformaticians in data analysis. The GUI design lowers the technical barrier for users without command-line experience while maintaining consistent parameter tracking and reproducibility. With a simple double-click of the JAR executable, users are presented with a main window where they can select from a collection of tools that operate within a thread-safe instance. This bypasses dependency installation and execution difficulties for users unfamiliar with command-line workflows. Each tool presents a unique window with a series of specific parameter selections. Tool parameter options are dynamically enabled or disabled based on the user’s selections to prevent invalid configurations. These features were designed to make the suite more intuitive for users and to reduce the learning curve required to get from starting an analysis to producing results.

While users may run ScriptManager on their local workstations, those interested in leveraging greater computing resources on their servers can still run the ScriptManager GUI by leveraging popular set-ups like X11 forwarding and Open OnDemand (OOD) [27]. By running ScriptManager on high-performance computing resources, users can increase threading options on more computationally intensive scripts to fully utilize available compute resources.

Each ScriptManager tool includes a command-line (CLI) equivalent, enabling execution on remote or headless systems. This allows users more comfortable with the command line or those who work on remote resources without support for graphical interactions to execute ScriptManager through its CLI. This framework also enables users with minimal command-line experience the opportunity to practice building scripts and workflows using the CLI for their analysis.

### Session logging

Analysis pipelines are often iteratively designed such that the final form of an analysis workflow is different from the first version. Bioinformaticians usually explore different methodologies or parameters before finalizing an analysis. As a result, at the time of publication, every workflow parameter must be documented precisely for the analysis to be reproducible. One challenge with GUI-based analysis is that to reproduce an analysis, a user must remember or record the exact parameter selections made during the GUI session if they wish to maintain reproducibility.

To facilitate reproducible analysis through the GUI, a logging feature is embedded in ScriptManager to track the exact workflow sequence performed during a GUI session. Upon execution of any script in the ScriptManager suite, a log record is initialized with a starting timestamp, an “incomplete” status, information on which script was executed, and all inputs and options passed to the script. All this information for each log record in a session can be viewed in the Logging Manager window’s “View Log” tab (**Fig. 1B**). The status updates when the script either fails or runs to completion, at which point the end timestamp is recorded. This produces a record that users can use to track the order and total runtimes of their workflow steps. For bulk processing or independent multi-file runs, the equivalent of several CLI execution statements for each input file is recorded in the log.

The information from a session’s log can be exported to a file as an executable shell script from the “Log Settings” tab in the Logging Manager (**Fig. 1C**). By saving a log for each session, the user can keep a record of the exact analysis used to produce a result or figure that can be included in their publication to support more rigorous and reproducible analysis. This tab also allows the user to toggle logging on/off, set the log verbosity, and change the destination file path. The saved log itself is a flat text file that can be directly executed as a shell script once a user updates the file path to the ScriptManager and Picard executable JAR files (**Fig. 1D**). These execution statements are a mapping of the GUI session’s script execution to the equivalent CLI execution so that GUI-based analyses can be re-run without manually clicking through the options and risking the introduction of errors.

In addition to keeping an exact record of an analysis, the saved log can facilitate collaboration between biochemists and bioinformaticians to design and develop new workflows. A biochemist analyzing their data might draft a workflow and export the saved log during the more dynamic development stages of a research project and then share their draft’s log with a bioinformatician. This log can serve as the exact specifications for building scalable, generalized workflows.

### Random seed control for stochastic methods

Many methods and approaches in bioinformatics are non-deterministic due to stochastic sampling of data. This means that the same user working on the same machine with the same installation configurations can re-execute an analysis method and get a distinct result. A common way of ensuring two different runs of the same tool give the same result in software development is to set a random seed. ScriptManager additionally supports user-specified random seeds for stochastic scripts in the suite. These tools support control and background analyses where reproducible seeds let readers and collaborators recapitulate the exact results of published analyses.

### Environment reproducibility and containerization

While shell scripts can capture the procedural record for a small set of samples in a single publication, scaling to hundreds or thousands of samples requires refactoring for parallel execution, error handling, and parameterization. For this, ScriptManager includes a Galaxy XML wrapper for each tool in the suite that can be run from the command line (available in the GitHub repository under ‘galaxy/’) to support interoperability with Galaxy, a popular pipeline engine [15]. Not only does this improve access to ScriptManager for users in the Galaxy community, but it also facilitates the incorporation of ScriptManager into more robust Galaxy workflows for standard, scalable, and reproducible data processing [28].

To meet the broader demand for portable, easy-to-install bioinformatics software, v1.0 also adds explicit support for two widely used reproducibility platforms: Singularity for containerized execution and Anaconda for environment management [19, 29]. A Singularity ‘.def’ under ‘singularity/’ is available to enable containerized execution that improves both reproducibility and cross-environment portability. Additionally, a bioconda recipe is now available for this release of ScriptManager.

### Unit/integration testing for version consistency

In software development, it is best practice to regularly run the software on test data, especially before a version release. This helps to ensure that errors are minimized during the development process of adding new features and fixing bugs. Following software development best practices, we built unit and integration tests that support ScriptManager’s continuous integration workflow. Regression testing provides cross-version output consistency, a key criterion for reproducibility in computational genomics workflows. This means for each future version of ScriptManager, a set of scripts is run to ensure that tools produce consistent outputs from given inputs across versions, minimizing the risk of regressions.

## Results

### TagPileup design and benchmarking resource scalability

To complement these reproducibility features, we benchmarked the platform’s most heavily used analytical module (TagPileup) across two axes: local single-machine performance and large-scale distributed execution. TagPileup calculates the occupancy of genomic reads across any set of genomic intervals (**Fig. 2A, 2B**). Given a BAM alignment file and a BED-formatted set of reference coordinates, TagPileup generates per-base-pair pileup matrices and average composite profiles that can be visualized as heatmaps and metaplots. A key design distinction is that TagPileup supports the selection of specific positions on each read fragment, such as the 5’ or 3’ end of either read in a paired-end dataset, or the midpoint of the inferred fragment, rather than tallying coverage across the entire read length (**Fig. 2C**). This preserves the base-pair resolution encoded by many genomic assays where the biological signal of interest is marked at a specific position on the DNA library fragment. TagPileup additionally provides filters for insert size, mate-pair status, and blacklisted regions, as well as strand-specific output, enabling users to interrogate specific subpopulations of library fragments. The practical application of these features across different assay types is demonstrated in the biological use cases below.

**Figure 2:**
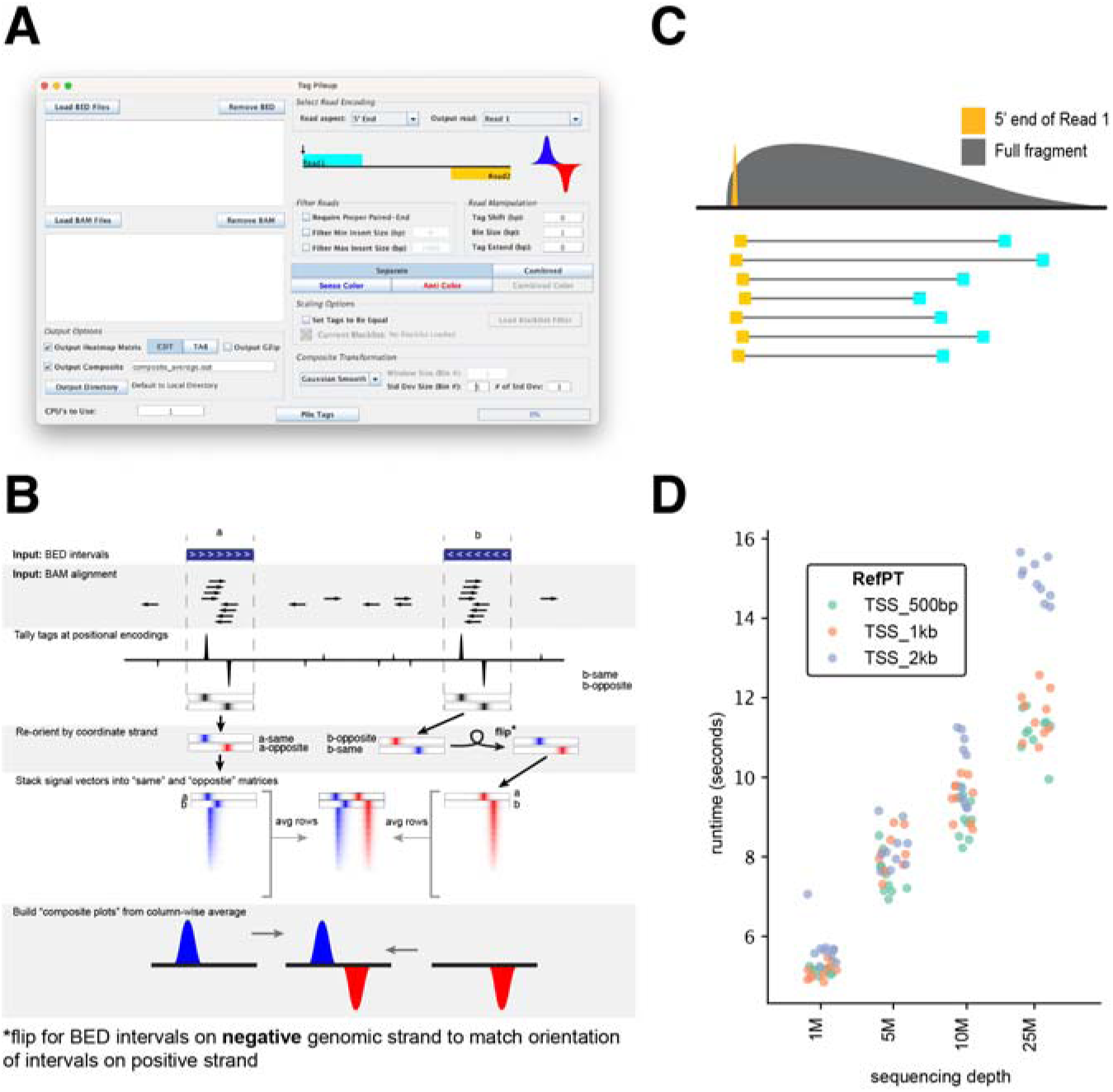
TagPileup is the high-resolution genomic analysis workhorse of ScriptManager. (A) TagPileup exposes a configurable pileup pipeline whose runtime depends on input size and the parameters chosen below; we benchmark the dominant cost factors in (D). (B) Schematic of TagPileup behavior on toy ChIP-exo data. The browser snippet shows two BED-coordinate intervals together with their BAM-aligned reads. Tags are tallied for each interval in a strand-aware manner, then transformed into per-interval vectors and aggregated into a tally matrix that can be visualized as a heatmap and averaged into a composite profile. (C) This pileup of toy bp-resolution data shows the profile of piling up tags from just the 5’ end of read 1 compared to the full fragment and how the full fragment choice can effectively smooth high-resolution signal. (D) TagPileups were performed on deeply sequenced nuclease-digested chromatin alignments subsampled to four different sequencing depths across three different window expansion lengths for 1,000 random TSS-centered reference points. Each TagPileup was repeated 10 times and timed to show how the TagPileup runtime scales with BAM file size and BED coordinate interval length.

To benchmark TagPileup performance, we down-sampled a deeply sequenced nuclease digestion protection dataset to 1M, 5M, 10M, and 25M paired-end reads and performed TagPileup ten times at each depth on 1,000 transcription start sites (TSS) expanded to three window lengths (500 bp, 1 kb, and 2 kb). All runs were allocated one CPU core and 16 GB of memory. Runtimes were comparable across window lengths at lower sequencing depths and diverged modestly at 25M paired-end reads (**Fig. 2D**), demonstrating that TagPileup scales efficiently for datasets typical of genomic experiments.

### ScriptManager is scalable with HPC– and HTC-style computing

To demonstrate ScriptManager’s scalability across heterogeneous computing platforms, we performed a large-scale High Throughput Computing (HTC) analysis using the containerized (Singularity) distribution of ScriptManager on the Open Science Grid (OSG) [30, 31]. Over 10,000 TagPileup jobs were executed across the Yeast Epigenome Project (YEP) dataset of ChIP-exo samples (1,145 BAM files) and nine reference point sets (TSS and eight motif BED files), submitted as a single HTCondor cluster of 10,305 jobs (**Fig. 3A; Additional File 2**). Jobs were dispatched across 36 distinct OSG sites in the United States (**Fig. 3B**). Nearly all jobs spent 90 minutes or less in the queue and completed within 2 hours (**Fig. 3C**). TagPileup execution itself completed in under two minutes per job, with runtimes scaling as a function of BAM file depth and the number of sites in the BED coordinate file (**Fig. 3D**). In aggregate, the workload consumed approximately 17.5 CPU-hours but completed in roughly two hours of wall-clock time, a ∼9x effective speedup over serial execution on a single core. These results illustrate that containerized TagPileup-based analyses can be deployed at consortium scale on shared HTC infrastructure, enabling large comparative-epigenomics studies without local HPC investment.

**Figure 3:**
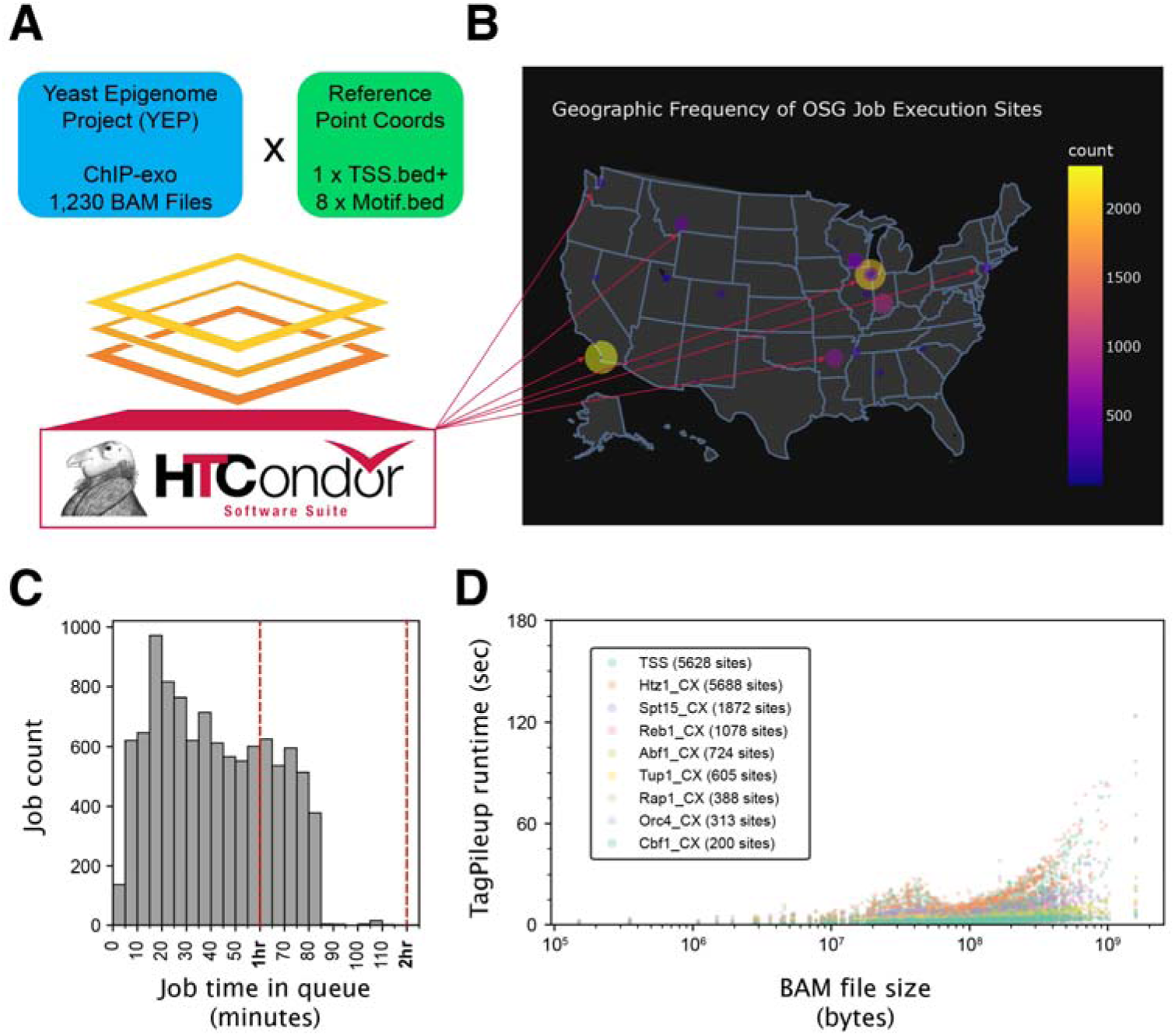
ScriptManager can be containerized and run on HTC systems. (A) Over 10,000 jobs, each performing a TagPileup of one YEP sample against one of nine reference points, were submitted to OSG using the HTCondor submission system. (B) The submitted jobs were scattered to computing resources across the USA for highly parallelized execution. (C) The distribution of time spent in the queue is visualized as a histogram while (D) the actual execution time of tag pileup is shown as a function of the sample dataset size and colored by the reference point used.

### Application Examples

Having established that TagPileup is performant and scalable, we now demonstrate how its parameter flexibility supports analysis across genomic assay types. Because different assays encode biological information at different positions within their DNA library fragments, the ability to select specific read encodings, apply fragment-level filters, and control strand output allows users to tailor their analysis to the structure of each dataset. We demonstrate this across several assay types, each motivating a different combination of TagPileup parameters (**Fig. 4; Additional File 3**).

**Figure 4:**
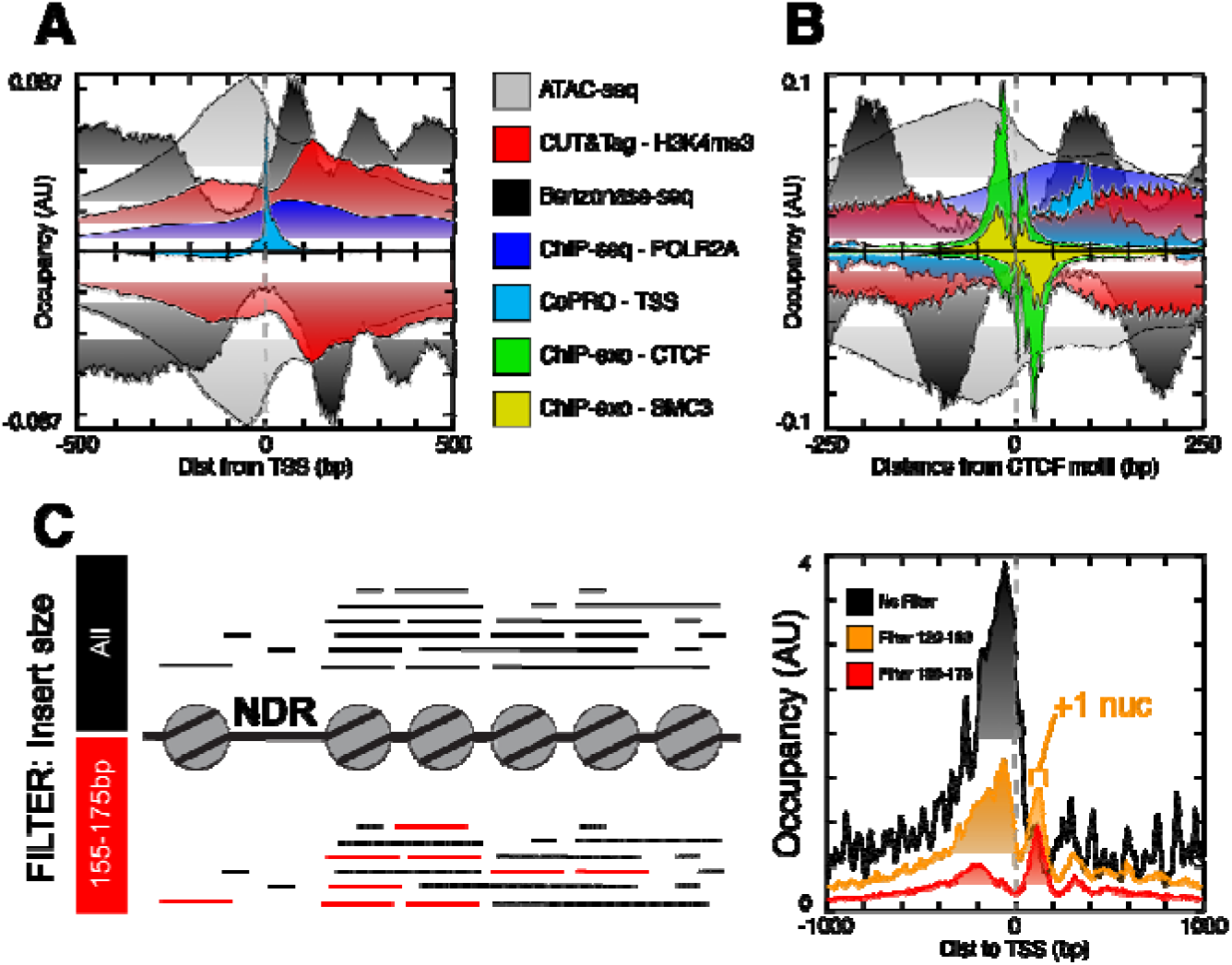
TagPileup supports diverse analysis of genomic data by leveraging information encoded in different parts of sequenced library fragments. Different assay types that encode information at distinct positions within the library fragment are visualized around two sets of reference points or genomic loci as an average or composite profile: **(A)** TSSs and **(B)** CTCF motif instances. CoPRO datasets were piled up at the 5’ end of read 2, which encodes the capped 5’ end of the nascent transcript (i.e., TSS or PRO-cap readout). RNA Pol II ChIP-seq datasets were piled up across the whole fragment alignment (5’ end of read 1 through 5’ end of read 2). Benzonase-seq was piled up at the cut sites on either end of each library fragment (just the 5’ end of read 1 and read 2) to mark unprotected and rotationally exposed DNA. ATAC-seq and CUT&Tag datasets were piled up at the inferred midpoint between paired reads, marking accessible chromatin regions (ATAC-seq) or accessible regions enriched for a specific target (CUT&Tag). For CTCF motif pileups **(B)**, two additional ChIP-exo datasets targeting CTCF and a cohesin subunit (SMC3) were piled up on their exonuclease cut sites to reveal crosslink-protected positions (5’ end of read 1). These composite profiles showcase TagPileup’s flexibility for interrogating different genomic features within and across different datasets. **(C)** Filtering DNase-seq fragments by insert size to select approximately mono-nucleosome-sized fragments (∼147 bp) and piling up the signal (library fragment midpoint) improves resolution of nucleosome phasing by enriching for fragments that span a single nucleosome and depleting shorter fragments that

Around annotated TSSs (UCSC knownCanonical), the pileups reveal distinct and complementary views of promoter architecture (**Fig. 4A**). CoPRO shows highly focused transcription initiation at the canonical TSS [32–36], while RNA Pol II ChIP-seq shows enrichment at the TSS with a gradual decline downstream [33–36], reflecting the transition from initiation to pausing and elongation. ATAC-seq [34] marks a region of open chromatin just upstream of the TSS, and nucleosome positioning from Benzonase-seq [37] and H3K4me3 CUT&Tag [38] reveals translational phasing beginning with the +1 nucleosome approximately 100 bp downstream.

The aligned sequence reads from these assays were then piled up around CTCF motif instances from the JASPAR database, along with two ChIP-exo datasets targeting CTCF and a cohesin subunit (SMC3) (**Fig. 4B**). This combination allows examination of the architectural organization of nucleosomes, TFs, and transcriptional activity around the contact sites of 3D chromatin loops. ChIP-exo, a high-resolution variant of ChIP-seq, requires selection of the 5’ end of read 1 to resolve precise crosslink positions [37, 39, 40]. The resulting profiles reveal the exact crosslink positions of CTCF at its motif, with SMC3 showing a matched crosslink pattern consistent with cohesin loading at CTCF-bound sites [37]. Nucleosomes appear roughly symmetrically phased around the motif (Benzonase-seq and CUT&Tag), whereas transcriptional activity is biased downstream (CoPRO and RNA Pol II ChIP-seq) and chromatin accessibility is biased upstream (ATAC-seq).

Beyond read encoding selection, TagPileup’s fragment-level filters enable users to examine specific subpopulations within a genomic library. For example, when working with paired-end nuclease protection data such as DNase-seq [41], applying an insert-size filter to retain only mono-nucleosome-sized fragments (∼147 bp) and piling up fragment midpoints enhances the nucleosome positioning signal, revealing sharper translational phasing by excluding sub-nucleosome– and multi-nucleosome-sized fragments (**Fig. 4C**).

These examples illustrate how the combinatorial flexibility of TagPileup’s read encoding and filters supports the analysis of diverse genomic assays within a single tool. Additional parameter combinations, assay-specific workflows, and features such as blacklist filtering are detailed as vignettes in the ScriptManager documentation.

## Documentation and Accessibility

### User-facing tutorials and vignettes

Comprehensive documentation underpins ScriptManager’s emphasis on accessibility and reproducibility. The platform’s online documentation provides step-by-step vignettes demonstrating how to analyze diverse genomic assays including ChIP-exo, ATAC-seq, RNA-seq, and CoPRO, using publicly available datasets. Each tutorial connects biological context with practical analysis steps, guiding users from raw data to interpretable visualizations.

Unlike canned analyses from published pipelines tailored to answering specific questions for certain genomic assays, ScriptManager supports a variety of ways for the user to analyze a given genomic dataset. ScriptManager has been used to analyze ChIP-exo, ChIP-seq, ATAC-seq, MNase-seq, RNA-seq, PIP-seq, PRO-seq/PRO-cap, and many other types of genomics datasets [37, 40, 42–50]. To demonstrate the use of ScriptManager to perform some standard genomics analysis, the ScriptManager documentation includes vignettes for some popular genomic assays under the “Examples” section, using publicly available datasets (**Table 1; Additional File 4**).

**Table 1:** List of available ScriptManager tutorial examples.

| Tutorial name | Purpose |
| --- | --- |
| ChIP-exo tutorial | Highlight utility of ScriptManager for building figures that showcase data with bp resolution and strand-specific properties |
| Genomic sequence analysis tutorial | Showcase visualization of annotation-based datasets by analyzing sequence content and positioning |
| Genome browser tracks tutorial | Users interact more directly with data through a genome browser by aggregating the dataset signal into a genome track file (BigWig) |
| ATAC-seq tutorial | Showcase data that is encoded across the entire read fragment |
| Co-PRO tutorial | Showcase data with information encoded at both ends of library fragments |
| **Table 1: List of available ScriptManager tutorial examples.** |  |

The ChIP-exo tutorial illustrates strand-specific read pileups and base-pair–resolution heatmaps, while the ATAC-seq example demonstrates strand-agnostic fragment-level profiling. Additional examples show how to construct BigWig genome browser tracks and perform motif-based sequence visualization. Together, these examples demonstrate how users can customize analyses by substituting datasets, modifying reference feature sets, or applying filters such as insert size or strand.

### Developer-facing API documentation

As an open-source suite of relatively independent scripts, ScriptManager’s structure is conducive to a community development approach, and its codebase includes useful functions for bioinformaticians who do Java-based development. To support a community of developers directly working with the growing ScriptManager codebase, ScriptManager needed developer-facing documentation beyond the existing user documentation. ScriptManager’s user documentation website now also hosts full API documentation (JavaDocs) for developers, ensuring transparency of internal methods and simplifying community contributions. By combining executable tutorials, reusable workflows, and open developer documentation, ScriptManager provides a comprehensive educational and technical foundation for FAIR-compliant, reproducible genomic data analysis.

## Discussion

ScriptManager occupies a distinct position in the genomic software landscape by combining the exploratory accessibility of a GUI-based tool with the reproducibility infrastructure typically associated with formal workflow systems. Established frameworks such as Galaxy and Nextflow provide powerful provenance tracking and scalable execution but require users to learn domain-specific workflow languages or web-based pipeline editors before they can begin analysis [15, 16]. ScriptManager’s GUI lowers this entry barrier, enabling wet-bench researchers to prototype analyses interactively, while the session logging feature automatically captures every parameter selection as an equivalent CLI command. The resulting exportable shell scripts serve as exact, executable specifications that can be handed to bioinformaticians for integration into more formal pipeline infrastructure. This GUI-to-CLI bridge addresses a practical gap that neither standalone GUI tools (which typically lack systematic provenance capture) nor workflow managers (which typically lack low-barrier exploratory interfaces) fill on their own. Combined with support for containerized execution via Singularity, installation through Anaconda and Bioconda, and Galaxy XML wrappers for each tool, ScriptManager allows a single analysis to move from exploratory prototyping to reproducible, scalable deployment without being rewritten in a different framework.

Among tools that share ScriptManager’s focus on read pileup and figure generation for high-resolution genomic assays, including deepTools [51], ngs.plot [52], and EnrichedHeatmap [53], ScriptManager is differentiated by a combination of features rather than any single capability (**Table 2**).

**Table 2:** Feature comparison of ScriptManager and similar tools.

| **Feature** | **ScriptManager** | **deepTools** | **ngs.plot** | **EnrichedHeatmap** |
| --- | --- | --- | --- | --- |
| Language/Runtime | Java | Python | R, perl, Python | R/Bioconductor |
| Interface | GUI and CLI | CLI | CLI (R scripts) | R programming API |
| Installation | JRE | Python libraries | R packages | BiocManager::install() |
| Galaxy support | Yes | Yes | Yes | None |
| Container support | singularity | Docker and Singularity | None | None |
| Active maintenance (as of 2026) | Yes | Yes | Last source-affecting commit 2017 | Yes (Bioconductor release 3.22) |

To our knowledge, ScriptManager is the only platform that provides both a GUI and a CLI from a single executable, lowering the barrier for users without command-line experience while preserving scriptable execution for batch and remote workflows. Its single-JAR distribution requires only a Java runtime, sidestepping the language-specific dependency management that Python– (deepTools), R– (EnrichedHeatmap), and mixed-language (ngs.plot) tools rely on. ScriptManager also supports the broadest combination of containerization (Singularity) and workflow integration (Galaxy) among these peer tools and remains under active development. These differences position ScriptManager as a low-barrier option for researchers who need to move fluidly between interactive exploration, scripted execution, and containerized large-scale runs without rewriting their analysis across frameworks.

Several directions for future development follow naturally from the current release. ScriptManager’s analytical scope currently centers on positional-encoding assays, those in which biological signal is marked by the position of reads or read endpoints on the genome. Extending support to assays that encode signal through nucleotide changes, such as bisulfite sequencing and metabolic labeling approaches [54–56], is a priority for upcoming releases. On the infrastructure side, the development of nf-core modules for ScriptManager tools would broaden interoperability with the Nextflow-based pipelines increasingly adopted at genomics centers. Expanded tutorials demonstrate how to integrate multiple assay types within a single workflow. For example, combining ChIP-exo occupancy profiles with ATAC-seq accessibility data at the same regulatory elements further demonstrate the cross-assay flexibility of the platform’s modular design. As an open-source project with full API documentation (JavaDocs), ScriptManager is structured to support community-driven development of new modules, and we encourage contributions from researchers working with assay types not yet represented in the current tool suite.

## Conclusions

ScriptManager v1.0 provides a unified, modular framework for the analysis and visualization of diverse genomic and epigenomic datasets. Three advances define this release. First, an expanded tool suite and standardized use of common genomic file formats enable cross-assay interoperability within a single platform. Second, a set of integrated reproducibility features including GUI session logging with exportable shell scripts, random seed control, unit and integration testing, and support for containerized (Singularity), Anaconda, and Galaxy-based execution lowers the barrier between exploratory analysis and formal, shareable workflows. Third, benchmarking demonstrates that the TagPileup module scales efficiently from local workstations to consortium-scale distributed execution: a 10,305-job analysis on the Open Science Grid consumed roughly 17.5 CPU-hours yet completed in under 2 hours of wall-clock time across 36 OSG sites. Together, these capabilities enable researchers across computational skill levels to build, document, and reproduce publication-quality genomic analyses with minimal configuration overhead.

## Availability of Source code and Requirements

- Project name: ScriptManager
- Project home page: https://github.com/CEGRcode/scriptmanager
- Operating system(s): Platform independent
- Programming language: Java
- Other requirements: Java 17 or higher
- License: MIT
- RRID: SCR_021797

## Additional Files

- **Additional File 1** – List of all scripts
- **Additional File 2** – Composite results from Figure 3 TagPileup runs on OSG-HTCondor
- **Additional File 3** – List of datasets used for Figure 4 and their accessions

## Declarations

**Ethics approval and consent to participate: not applicable**

**Consent for Publication: not applicable**

## Availability of data and materials

Source for ScriptManager is available at https://github.com/CEGRcode/scriptmanager. The public datasets analyzed for this manuscript include data from ENCODE [33–36] and several publicly available Gene Expression Omnibus (GEO) projects: GSE147927 [46], GSE266494 [37], GSE267711 [37], and GSE124557 [38]. Exact accessions available in **Additional File 3**. Scripts used to generate the figures are available on GitHub under the “paper-v1.0” subdirectory (https://github.com/CEGRcode/scriptmanager-docs) and the full documentation website is hosted at https://pughlab.mbg.cornell.edu/scriptmanager-docs/.

## Author contributions

OWL contributed to and rearchitected the codebase, ran performance metrics, supervised development, and co-wrote the manuscript. BB implemented new scripts, added gzip support, and drafted the technical documentation. CL and DZ contributed to implementing Galaxy support and testing. AD contributed to tutorials and user documentation for the website. BFP provided biochemical and genomics insight (design), suggested useful utilities to include, and provided feedback on the manuscript. WKML conceptualized the project, contributed to the codebase, provided project guidance and oversight, and co-wrote the manuscript.

## List of Abbreviations

CDT: Clustered Data Table file format
CLI: Command Line Interface
GUI: Graphical User Interface
HPC: High Performance Computing
HTC: High Throughput Computing
JAR: Java ARchive format
JRE: Java Runtime Environment
OOD: Open OnDemand
OSG: Open Science Grid
TSS: Transcription start site
TSV: Tab Separated Values file format
TXT: Text file format
YEP: Yeast Epigenome Project

## Acknowledgements

We thank the members of the Pugh Lab and members of Cornell University Biotechnology Resource Center Epigenomics Core Facility for their help with user testing and suggestions for improving the UI/UX of the interface with a special thanks to Abeer Almutairy for her contributions to the peak-align tool. Sample processing and FAIR-compliant analysis made possible by Platform for Epigenomic and Genomic Research (RRID:SCR_021861). Computational support and services provided by Penn State Institute for Computational and Data Sciences (RRID:SCR_025154), Penn State Center for Applications of Artificial Intelligence and Machine Learning to Industry (RRID:SCR_022867), and the OSG Consortium, which is supported by National Science Foundation awards #2030508 and #2323298 [30, 31, 57, 58]. This work also used Jetstream2 at Indiana University through allocation BIO250427 (OW Lang) from the Advanced Cyberinfrastructure Coordination Ecosystem: Services & Support (ACCESS) program, which is supported by National Science Foundation grants #2138259, #2138286, #2138307, #2137603, and #2138296. We also acknowledge the following resources: Picard (RRID:SCR_006525); Galaxy (RRID:SCR_006281); GenoPlotter (RRID: SCR_028339); Cornell University Biotechnology Resource Center Epigenomics Core Facility (RRID:SCR_021287).

## References

1. Policastro RA, Zentner GE: Global approaches for profiling transcription initiation. Cell Rep Methods 2021, 1(5).

2. Satam H, Joshi K, Mangrolia U, Waghoo S, Zaidi G, Rawool S, Thakare RP, Banday S, Mishra AK, Das G et al: Next-Generation Sequencing Technology: Current Trends and Advancements. Biology (Basel) 2023, 12(7).

3. Andrews S: FastQC: a quality control tool for high throughput sequence data. In.; 2010.

4. Bonfield JK, Marshall J, Danecek P, Li H, Ohan V, Whitwham A, Keane T, Davies RM: HTSlib: C library for reading/writing high-throughput sequencing data. Gigascience 2021, 10(2).

5. Quinlan AR, Hall IM: BEDTools: a flexible suite of utilities for comparing genomic features. Bioinformatics 2010, 26(6):841–842.

6. Li H, Durbin R: Fast and accurate short read alignment with Burrows-Wheeler transform. Bioinformatics 2009, 25(14):1754–1760.

7. Lai WK, Bard JE, Buck MJ: ArchTEx: accurate extraction and visualization of next-generation sequence data. Bioinformatics 2012, 28(7):1021–1023.

8. Neumann T, Herzog VA, Muhar M, von Haeseler A, Zuber J, Ameres SL, Rescheneder P: Quantification of experimentally induced nucleotide conversions in high-throughput sequencing datasets. BMC Bioinformatics 2019, 20(1):258.

9. Gordon A: FASTQ/A short-reads pre-processing tools. In.; 2009.

10. Ziemann M, Poulain P, Bora A: The five pillars of computational reproducibility: bioinformatics and beyond. Brief Bioinform 2023, 24(6).

11. Wilkinson MD, Dumontier M, Aalbersberg IJ, Appleton G, Axton M, Baak A, Blomberg N, Boiten JW, da Silva Santos LB, Bourne PE et al: The FAIR Guiding Principles for scientific data management and stewardship. Sci Data 2016, 3:160018.

12. Kim YM, Poline JB, Dumas G: Experimenting with reproducibility: a case study of robustness in bioinformatics. Gigascience 2018, 7(7).

13. Garijo D, Kinnings S, Xie L, Xie L, Zhang Y, Bourne PE, Gil Y: Quantifying reproducibility in computational biology: the case of the tuberculosis drugome. PLoS One 2013, 8(11):e80278.

14. Mangul S, Mosqueiro T, Abdill RJ, Duong D, Mitchell K, Sarwal V, Hill B, Brito J, Littman RJ, Statz B et al: Challenges and recommendations to improve the installability and archival stability of omics computational tools. PLoS Biol 2019, 17(6):e3000333.

15. Galaxy Community: The Galaxy platform for accessible, reproducible, and collaborative data analyses: 2024 update. Nucleic Acids Res 2024, 52(W1):W83–W94.

16. Di Tommaso P, Chatzou M, Floden EW, Barja PP, Palumbo E, Notredame C: Nextflow enables reproducible computational workflows. Nat Biotechnol 2017, 35(4):316–319.

17. Deelman E, Vahi K, Rynge M, Mayani R, da Silva RF, Papadimitriou G, Livny M: The Evolution of the Pegasus Workflow Management Software. Computing in Science & Engineering 2019, 21(4):22–36.

18. Anaconda Software Distribution. In.; 2016.

19. Kurtzer GM, Sochat V, Bauer MW: Singularity: Scientific containers for mobility of compute. PLoS One 2017, 12(5):e0177459.

20. Merkel D: Docker: lightweight Linux containers for consistent development and deployment. Linux J 2014, 2014(239).

21. National Science Foundation: Preparing Your Data Management and Sharing Plan. In.

22. National Institutes of Health: Data Management & Sharing Policy Overview. In.; 2023.

23. Simoneau J, Dumontier S, Gosselin R, Scott MS: Current RNA-seq methodology reporting limits reproducibility. Brief Bioinform 2021, 22(1):140–145.

24. Nekrutenko A, Taylor J: Next-generation sequencing data interpretation: enhancing reproducibility and accessibility. Nat Rev Genet 2012, 13(9):667–672.

25. Lang O, Pugh BF, Lai WKM: ScriptManager: an interactive platform for reducing barriers to genomics analysis. In: Practice and Experience in Advanced Research Computing; Boston, MA, USA. Association for Computing Machinery 2022: Article 33.

26. Picard: Picard toolkit. In: Broad Institute, GitHub repository, https://broadinstitute.github.io/picard/. 2019.

27. Hudak DE, Bitterman T, Carey P, Johnson D, Franz E, Brady S, Diwan P: OSC OnDemand: a web platform integrating access to HPC systems, web and VNC applications. In.: 2013.

28. Blankenberg D, Von Kuster G, Bouvier E, Baker D, Afgan E, Stoler N, Galaxy T, Taylor J, Nekrutenko A: Dissemination of scientific software with Galaxy ToolShed. Genome Biol 2014, 15(2):403.

29. Gruning B, Dale R, Sjodin A, Chapman BA, Rowe J, Tomkins-Tinch CH, Valieris R, Koster J, Bioconda T: Bioconda: sustainable and comprehensive software distribution for the life sciences. Nat Methods 2018, 15(7):475–476.

30. Pordes R, Petravick D, Kramer B, Olson D, Livny M, Roy A, Avery P, Blackburn K, Wenaus T, Würthwein F et al: The open science grid. J Phys Conf Ser 2007, 78:012057.

31. OSG: OSPool. OSG 2006.

32. Tome JM, Tippens ND, Lis JT: Single-molecule nascent RNA sequencing identifies regulatory domain architecture at promoters and enhancers. Nat Genet 2018, 50(11):1533–1541.

33. Consortium EP: An integrated encyclopedia of DNA elements in the human genome. Nature 2012, 489(7414):57–74.

34. Kagda MS, Lam B, Litton C, Small C, Sloan CA, Spragins E, Tanaka F, Whaling I, Gabdank I, Youngworth I et al: Data navigation on the ENCODE portal. Nat Commun 2025, 16(1):9592.

35. Zhang J, Lee D, Dhiman V, Jiang P, Xu J, McGillivray P, Yang H, Liu J, Meyerson W, Clarke D et al: An integrative ENCODE resource for cancer genomics. Nat Commun 2020, 11(1):3696.

36. Hitz BC, Lee J-W, Jolanki O, Kagda MS, Graham K, Sud P, Gabdank I, Seth Strattan J, Sloan CA, Dreszer T et al: The ENCODE Uniform Analysis Pipelines. bioRxiv 2023:2023.2004.2004.535623.

37. Chen H, Krebs JE, Lang OW, Hu J, Melini D, Hyle J, Li C, Lai WKM, Pugh BF: Genome-wide rotational and translational phasing of nucleosomes with human transcription factors. Mol Cell 2026.

38. Kaya-Okur HS, Wu SJ, Codomo CA, Pledger ES, Bryson TD, Henikoff JG, Ahmad K, Henikoff S: CUT&Tag for efficient epigenomic profiling of small samples and single cells. Nat Commun 2019, 10(1):1930.

39. Rhee HS, Pugh BF: Genome-wide structure and organization of eukaryotic pre-initiation complexes. Nature 2012, 483(7389):295–301.

40. Rossi MJ, Lai WKM, Pugh BF: Simplified ChIP-exo assays. Nat Commun 2018, 9(1):2842.

41. Crawford GE, Holt IE, Whittle J, Webb BD, Tai D, Davis S, Margulies EH, Chen Y, Bernat JA, Ginsburg D et al: Genome-wide mapping of DNase hypersensitive sites using massively parallel signature sequencing (MPSS). Genome Res 2006, 16(1):123–131.

42. Lai WK, Pugh BF: Genome-wide uniformity of human ’open’ pre-initiation complexes. Genome Res 2017, 27(1):15–26.

43. Rossi MJ, Lai WKM, Pugh BF: Genome-wide determinants of sequence-specific DNA binding of general regulatory factors. Genome Res 2018, 28(4):497–508.

44. Badjatia N, Rossi MJ, Bataille AR, Mittal C, Lai WKM, Pugh BF: Acute stress drives global repression through two independent RNA polymerase II stalling events in Saccharomyces. Cell Rep 2021, 34(3):108640.

45. Lai WKM, Mariani L, Rothschild G, Smith ER, Venters BJ, Blanda TR, Kuntala PK, Bocklund K, Mairose J, Dweikat SN et al: A ChIP-exo screen of 887 Protein Capture Reagents Program transcription factor antibodies in human cells. Genome Res 2021, 31(9):1663–1679.

46. Rossi MJ, Kuntala PK, Lai WKM, Yamada N, Badjatia N, Mittal C, Kuzu G, Bocklund K, Farrell NP, Blanda TR et al: A high-resolution protein architecture of the budding yeast genome. Nature 2021, 592(7853):309–314.

47. Mittal C, Lang O, Lai WKM, Pugh BF: An integrated SAGA and TFIID PIC assembly pathway selective for poised and induced promoters. Genes Dev 2022, 36(17-18):985–1001.

48. van Breugel ME, van Kruijsbergen I, Mittal C, Lieftink C, Brouwer I, van den Brand T, Kluin RJC, Hoekman L, Menezes RX, van Welsem T et al: Locus-specific proteome decoding reveals Fpt1 as a chromatin-associated negative regulator of RNA polymerase III assembly. Mol Cell 2023, 83(23):4205–4221 e4209.

49. Louder RK, Park G, Ye Z, Cha JS, Gardner AM, Lei Q, Ranjan A, Hollmuller E, Stengel F, Pugh BF et al: Molecular basis of global promoter sensing and nucleosome capture by the SWR1 chromatin remodeler. Cell 2024, 187(24):6849–6864 e6818.

50. Ranjan A, Elalaoui E, Tang X, Cha J, Louder RK, Nguyen K, Ye J, Bennani M, Gardner AM, Liu D et al: Histone acetylation readers Bdf1 and Yaf9 direct SWR1 remodeler to +1 nucleosome. Sci Adv 2025, 11(32):eadt2002.

51. Ramirez F, Ryan DP, Gruning B, Bhardwaj V, Kilpert F, Richter AS, Heyne S, Dundar F, Manke T: deepTools2: a next generation web server for deep-sequencing data analysis. Nucleic Acids Res 2016, 44(W1):W160–165.

52. Shen L, Shao N, Liu X, Nestler E: ngs.plot: Quick mining and visualization of next-generation sequencing data by integrating genomic databases. BMC Genomics 2014, 15:284.

53. Gu Z, Eils R, Schlesner M, Ishaque N: EnrichedHeatmap: an R/Bioconductor package for comprehensive visualization of genomic signal associations. BMC Genomics 2018, 19(1):234.

54. Herzog VA, Reichholf B, Neumann T, Rescheneder P, Bhat P, Burkard TR, Wlotzka W, von Haeseler A, Zuber J, Ameres SL: Thiol-linked alkylation of RNA to assess expression dynamics. Nat Methods 2017, 14(12):1198–1204.

55. Gu H, Smith ZD, Bock C, Boyle P, Gnirke A, Meissner A: Preparation of reduced representation bisulfite sequencing libraries for genome-scale DNA methylation profiling. Nat Protoc 2011, 6(4):468–481.

56. Feil R, Charlton J, Bird AP, Walter J, Reik W: Methylation analysis on individual chromosomes: improved protocol for bisulphite genomic sequencing. Nucleic Acids Res 1994, 22(4):695–696.

57. OSG: Open Science Data Federation. OSG 2015.

58. Sfiligoi I, Bradley DC, Holzman B, Mhashilkar P, Padhi S, Wurthwein F: The pilot way to grid resources using glideinWMS. 2009 WRI World Congress on Computer Science and Information Engineering 2009, 2:428–432.

